# Condensin II is required for efficient Spindle Assembly Checkpoint activation in Drosophila male meiosis

**DOI:** 10.1101/2022.02.23.481572

**Authors:** Cintia Horta, Alexandra Tavares, Raquel A. Oliveira

## Abstract

Reductional nuclear division in meiosis is essential for diploid life. A fundamental event in meiosis is chromatin condensation, through mechanisms not yet fully understood. Current data suggest that Condensins are key players in building and sustaining mitotic and meiotic chromosome structure. In *Drosophila*, Condensin II appears to be dispensable for faithful mitosis in somatic tissues yet essential in the germline. Previous work has demonstrated that in *Drosophila* male meiosis, Condensin II is required for the segregation of homologous chromosomes into distinct territories during prophase I, possibly through the resolution of chromosomal intertwines. Here we show that in addition to this well-established function in meiotic chromatin assembly, Condensin II is required for robust Spindle Assembly Checkpoint (SAC) signaling in male meiosis. In the absence of Condensin II, spermatocytes undergo faster meiotic divisions and display reduced ability to prolong meiosis in the presence of spindle poisons. This is attributed to the inability to recruit a key SAC component (Mad1) to the kinetochore. Importantly, we demonstrate that the absence of a robust SAC response in Condensin II mutants, and consequent accelerated meiosis, is a strong contributor to the meiotic defects associated with these mutants. We show that artificial prolongation of meiotic divisions, using conditions that delay anaphase onset in a SAC-independent manner, is sufficient to rescue segregation defects and aneuploidy associated with Condensin II mutations. We therefore conclude that Condensin II can be dispensable for the resolution of topological problems and chromosome condensation if cells are able to prolong meiosis. Yet, the newly found role of this complex in the robustness of the SAC reduces meiotic timing leading to severe chromosome segregation defects.

## Introduction

Proper chromosome assembly is critical for the faithful division of genetic material, both in mitosis and meiosis. Although the mechanisms underlying chromosome compaction are not yet fully understood, it is established that condensin complexes are major players in the process. Condensin complexes are composed by two structural maintenance of chromosome (SMC) proteins (SMC2 and SMC4) that are linked by a kleisin subunit (Barren/Cap-H for Condensin I and Barren2/Cap-H2 for condensin II) [1, 2]. These complexes also contain HEAT-repeats proteins (Cap-G/G2 and Cap-D2/D3) thought to play regulatory roles [3]. The exact mechanisms by which Condensins regulate DNA morphology are still unknown, but research in recent years support that condensins are responsible for organizing mitotic chromosomes through the ATP-dependent extrusion of DNA loops [4, 5]. Condensin’s activity ensures not only chromosome compaction but also their structural integrity, by promoting the correct physical properties (e.g. rigidity) [6–9], and the efficient resolution of chromosome intertwines, possibly by guiding the activity of topoisomerase 2 [10, 11].

The presence of two distinct complexes in several organisms raises some questions regarding their specialized functions and potential redundancies across the tree of life. Whereas metazoans possess two condensin complexes, yeast cells rely on the presence of a single condensin [2]. Moreover, the ratio between condensin I and II is very variable across species and impacts on chromosomal architecture in both interphase and mitosis [12, 13]. An extreme case in asymmetric condensin I/II usage is *Drosophila melanogaster*, where somatic mitosis relies mostly on condensin I [14, 15]. Condensin II, in contrast, is kept at very reduced amounts by SCF-Slimb ubiquitin ligase-induced degradation of CAP-H2 [16, 17], and is required mostly in the germline [18, 19].

The role of both condensin complexes has been mostly studied in mitotic divisions. Yet, several studies support that these are also critical for the assembly of meiotic chromosomes [20]. Condensin I or II deficiency in meiosis has been associated with abnormal chromosome morphology and chromatin bridges, sustaining that their primarily roles in meiosis are similar to the mitotic ones, i.e. chromosome compaction, rigidity and sister chromatid resolution. These functions are highly conserved across species, including yeast [21], mouse [9, 22], Drosophila [23], C. elegans [24, 25], plants [26, 27] and even the evolutionarily distant protist Tetrahymena, with non-canonical chromosome configurations [28]. Additionally, condensin complexes have been also reported to display some meiosis-specific functions, suggesting that these complexes may have acquired specific functions to fulfill unique meiotic events. For example, condensins have been proposed to contribute to multiple stages of meiotic recombination, including Synaptonemal Complex assembly, Double Strand Break (DSB) formation, DSB processing, number and distribution of cross-overs and the resolution of recombination-dependent links [29–32]. The role of condensin complexes in homologue pairing has been reported even in organisms without recombination, such as in *Drosophila* male meiosis I. In this organism, homologous chromosome pairing takes place during prophase I, when bivalents “segregate” to distinct nuclear regions, forming the so-called Chromosome Territories (CT) [33]. The mechanisms that drive this process are not fully understood but CT formation depends on condensin II activity [18, 34]. Mutants for this complex fail to display distinct CTs, possibly through the inability of resolving intra-chromosome intertwines, resulting in severe segregation defects [18].

Another unique event in meiotic divisions is the co-orientation of sister kinetochores in the first division. In yeast, condensin has been shown to directly contribute to this co-orientation process, thereby ensuring monopolar attachment of sister kinetochore that allows homologue segregation in meiosis I [35, 36]. Additionally, meiosis requires that cohesin, the molecular glue that holds chromosomes together, is not fully released in the first round of nuclear division [37]. Studies in yeast have suggest that condensin promotes cohesin release in meiosis [21]. In sharp contrast, condensin I has been recently proposed to contribute to cohesin protection in meiosis I in C. elegans [38]. Thus, the chromosomal landscape modulated by condensins and/or condensins’ additional functions ensure specific meiotic events, with potential differences across evolution.

Condensin complexes, particularly condensin II, have also been proposed to contribute to the assembly of specialized chromosomal regions as, for example, the centromeres [39–41]. Centromeres are critical structures for the fidelity of nuclear division as they serve as a platform for the assembly of the kinetochore, a proteinaceous structure that mediates chromosome attachment to the spindle. In addition to chromosome attachments, the kinetochore also catalyzes the production of the Mitotic Checkpoint Complex (MMC), and thereby ensures the activity of the Spindle Assembly Checkpoint (SAC), a surveillance mechanism that delays the onset of anaphase until all chromosomes are properly aligned [42]. Yet, most studies report that cells with compromised condensin activity are able to mount a robust SAC response, as evidenced by the fact that condensin depletion is often associated with a delay in mitotic or meiotic progression [6, 9, 43–45]. SAC activation is possibly caused by an indirect effect that chromosome assembly defects may have on the stability of chromosome attachment errors and/or defects in intra-kinetochore stretching required to silence the SAC [6, 7, 45]. In contrast with these observations, here we report that Condensin II mutations in *Drosophila* male meiosis do not delay meiotic progression but, on the contrary, accelerate meiosis timing. We reveal that in addition to its canonical role in chromosome resolution and CT formation, Condensin II is required for robust SAC response in this organism.

## Results

### Condensin II mutants undergo accelerated meiotic divisions

Mutations in condensin II subunits have been previously associated with failures to resolve chromosome entanglements in male germ cell divisions, characterized by defects in resolving homologues in different territories and severe chromosome bridges in both meiotic divisions [18]. We therefore aimed to investigate how the meiotic machinery would sense these topological problems. To address this question, we optimised live cell imaging approaches of *Drosophila* testis to gain a dynamic characterization of the meiotic process. We established a strain that was mutant for the Condensin II-specific subunit Cap-H2, *Cap-H2*^*e03210*^ (a null allele from the Drosophila Exilixis collection, see Fig. S1) over a deficiency for the same chromosomal region Df(3R)BSC529 (refered to as Cap-H2 mutant (Cap-H2^mut^) hereafter). Flies also expressed fluorescent labelled histones/tubulin to monitor meiotic progression. Quantitative analysis of meiotic timing was accessed by measuring the time from NEBD to the anaphase onset. The NEBD was estimated using HisH2Av-RFP, and defined by the first frame in which the nucleoplasm pool is dispersed showing a higher contrast of the chromatin. Anaphase onset was considered as the first frame in which chromatin is pulled to opposite poles with concomitant cell elongation. We anticipated that given the severe chromosome structural defects present in this mutant, and how these would likely impact on chromosome attachments to the spindle, cells would sense these defects and activate the Spindle Assembly Checkpoint response. In sharp contrast with this expectation, our analysis showed that in Cap-H2 mutants, meiotic divisions were happening significantly faster than in control strains. This was true for meiosis I (Figure 1A), but also meiosis II (Figure 1B). We determined that Cap-H2 mutants present a significant acceleration of approximately 10 minutes in meiosis I and 7 minutes in meiosis II, which represents ~30% decrease in the nuclear division timing. To estimate the degree of acceleration observed in Cap-H2 mutants, we compared the meiotic timings observed in this condition with the one present in a SAC-null situation [46]. In accordance with previous reports [47], we found that mad2 mutant flies - *mad2*^*P*^ - undergo accelerated meiotic divisions I (Figure 1 A and B). These results support that in contrast to previous reports [48], which argued that SAC is not functional in *Drosophila* male meiosis, this checkpoint controls the timing of male meiotic divisions. Most importantly, the division timing observed in Cap-H2 mutant was similar to the observed in flies without a functional SAC.

**Figure 1.**
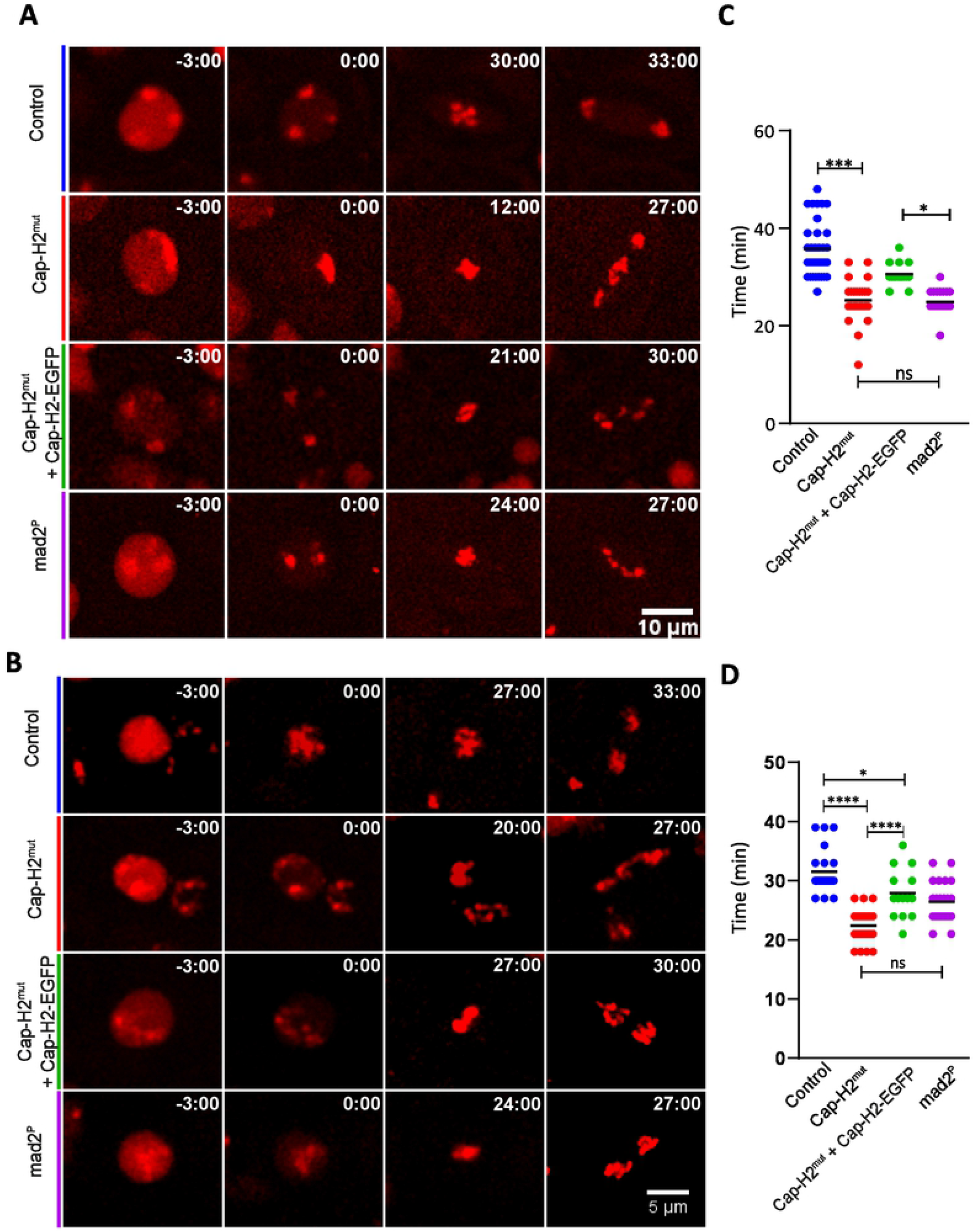
CAP-H2 mutant undergoes a faster meiosis. (**A, B**) Stills from live imaging of Control, Cap-H2^mut^, Cap-H2^mut^ + Cap-H2 rescue and mad2^P^ mutant spermatocytes undergoing meiosis I (A) and II (B). Time (min) is relative to NEBD. (**C, D**) Quantification of Meiosis I (C) and II (D) duration (from NEBD to anaphase onset) in all studied strains. Each dot represents an individual spermatocyte from at least 4 independent movies. *, P < 0.05; **, P < 0.01; ***, P < 0.001; ****, P < 0.0001 using ONE-way ANOVA (“Tukey’s multiple comparisons test”). Black lines indicate Median. Detailed genotypes can be found in Table S3.

To confirm that the obtained results were indeed attributed to Cap-H2 mutations, we performed rescue experiments. For this, we used a transgene that ectopically expresses Cap-H2 (Cap-H2^TEVA^-EGFP) which is able to rescue male fertility (Fig. S1) and nurse cell chromatin dispersal (not shown). It should be noted that this construct does not completely fulfil the endogenous Cap-H2 protein functions as CT formation is only partially restored (Fig. S1). Despite this, adding back a single copy of the Cap-H2 transgene to the mutant background was sufficient to partially restore the cell division timing, demonstrating that the observed acceleration in meiotic progression was Cap-H2-dependent (Figure 1). To test whether the faster meiotic timings are a general phenotype of condensin II, we performed similar analysis using mutants for another condensin II subunit (Cap-D3) [18], which displayed similar acceleration in meiotic timings (Fig. S2). These results led us to hypothesize that absence of condensin II results in defective Spindle Assembly Checkpoint response in both meiotic divisions.

### Cap-H2 Mutant has a compromised SAC response

We next aimed to address if accelerated meiotic timings are caused by an intrinsic reduced SAC robustness or, alternatively, are caused by modulation of microtubule-kinetochore attachments. Prior studies in *Xenopus* egg extracts or human tissue cells reported severe defects in kinetochore-microtubule interactions after Condensin depletion [49]. Interestingly, depletion of Condensins lead to an increase in merotelic attachments (when both kinetochores are attached by the same spindle pole) [41]. This kind of attachment is known to be undetected by the SAC which could account for a premature SAC satisfaction. To evaluate SAC competency independently of kinetochore-microtubule attachments, we tested if Cap-H2 mutants can establish a robust checkpoint response, and consequently delay meiotic exit when in the presence of microtubule depolymerizing drugs. Isolated testes were treated with 100 μM of colchicine to induce microtubule depolymerization, and live cell imaging was used to estimate the time of arrest (from NEBD to meiotic exit) in both meiotic divisions (Figure 2). Our results indicate that Cap-H2 mutant has a weaker checkpoint response when compared with control flies. Upon colchicine treatment, Cap-H2 mutant cells were able to delay meiotic exit for solely ~75 ± 18 minutes while control flies arrested 186 ± 24 minutes (Figure 2) in meiosis I. The same tendency is observed in the second meiotic division, where Cap-H2 mutants present a delay of 75 ± 6 min compared to the 104 ± 8 minutes arrest observed for control flies (Figure 2). Although displaying a compromised SAC response, Cap-H2 mutant was not behaving as a SAC null since the time of meiotic arrest was significantly higher than the observed for *mad2*^*P*^ mutant flies, which present the same division timing with and without the presence of microtubule poisons (Figure 2). These experiments revealed that condensin II mutants have a reduced ability to mount a robust SAC signalling, independently of the state of microtubule kinetochore attachments.

**Figure 2.**
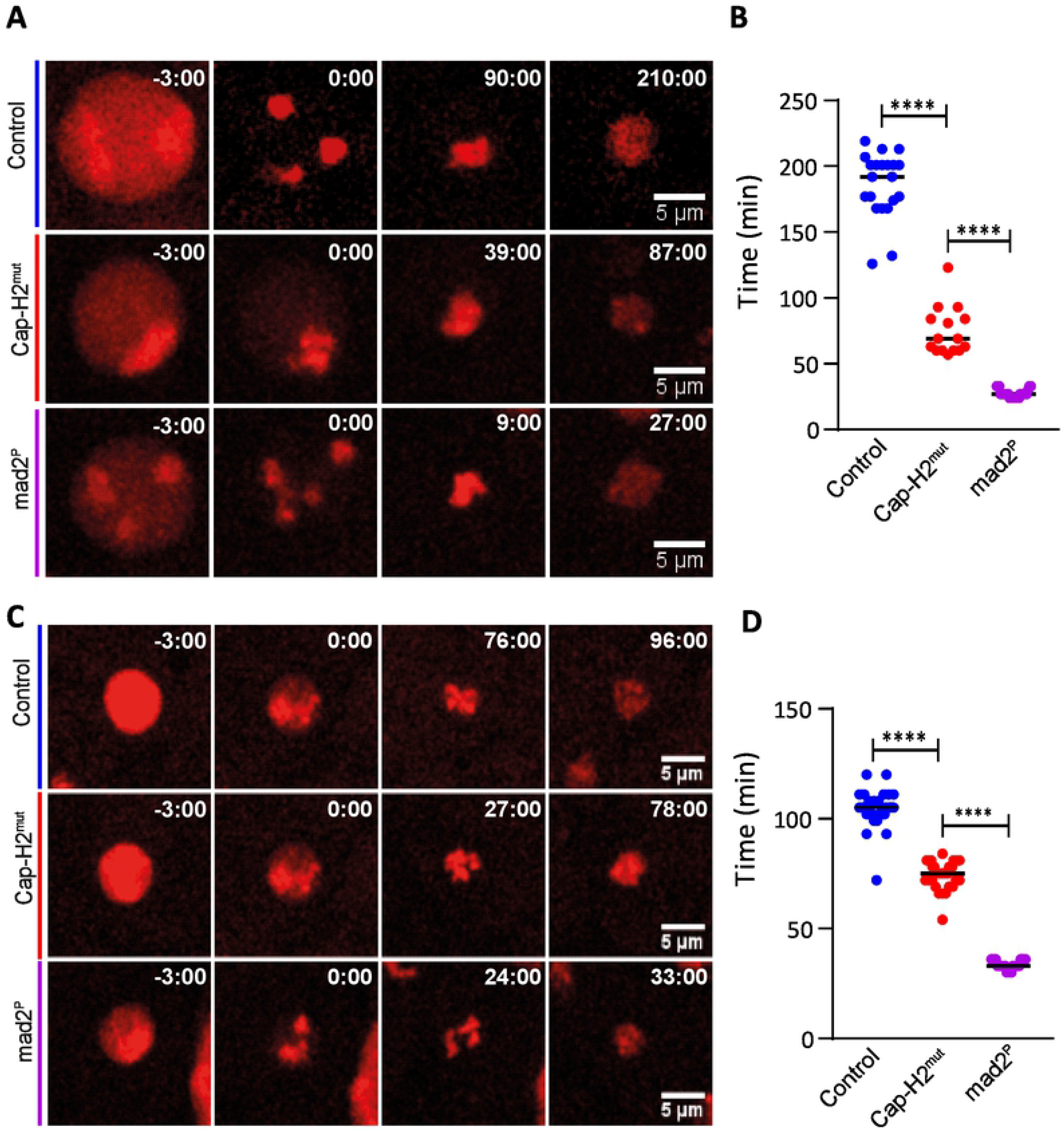
CAP-H2 mutant has a compromised SAC response. (**A-C**) Stills from live imaging of Control, Cap-H2^mut^, and mad2^P^ mutant spermatocytes undergoing meiosis I (A) or meiosis II (C) in the presence of 100 μM colchicine; Time (min) is relative to NEBD; scale bars are 5 μM (**B, D**) Nuclear division timing upon microtubule depolymerisation (100 μM colchicine), defined from NEBD and the time of chromosome decondensation. *, P < 0.05; **, P < 0.01; ***, P < 0.001; ****, P < 0.0001 using ONE-way ANOVA (“Tukey’s multiple comparisons test”). Black lines indicate Median. Detailed genotypes can be found in Table S3.

### Cap-H2 Mutants are defective at Mad 1 kinetochore recruitment

Our results so far suggest that Cap-H2 mutants are not fully checkpoint competent. In order to build a checkpoint response, SAC components must localize to unattached kinetochores. To test the ability of recruitment of SAC components in the absence of Cap-H2, we performed immunohistochemistry assays to access the levels of a key SAC component - Mad1 - upon microtubules depolymerization. We used quantitative imaging analysis to probe for the levels of Mad1 present at kinetochores. This analysis revealed that Cap-H2 mutants display a reduced ability for localizing Mad1 to the kinetochore upon colchicine arrest, in both meiotic divisions (Figure 3), being even more pronounced in meiosis II (Figure 3B). As expected, the levels of kinetochore-bound Mad1 are partially recovered by adding an exogenous copy of Cap-H2 to the mutant background (Figure 3).

**Figure 3.**
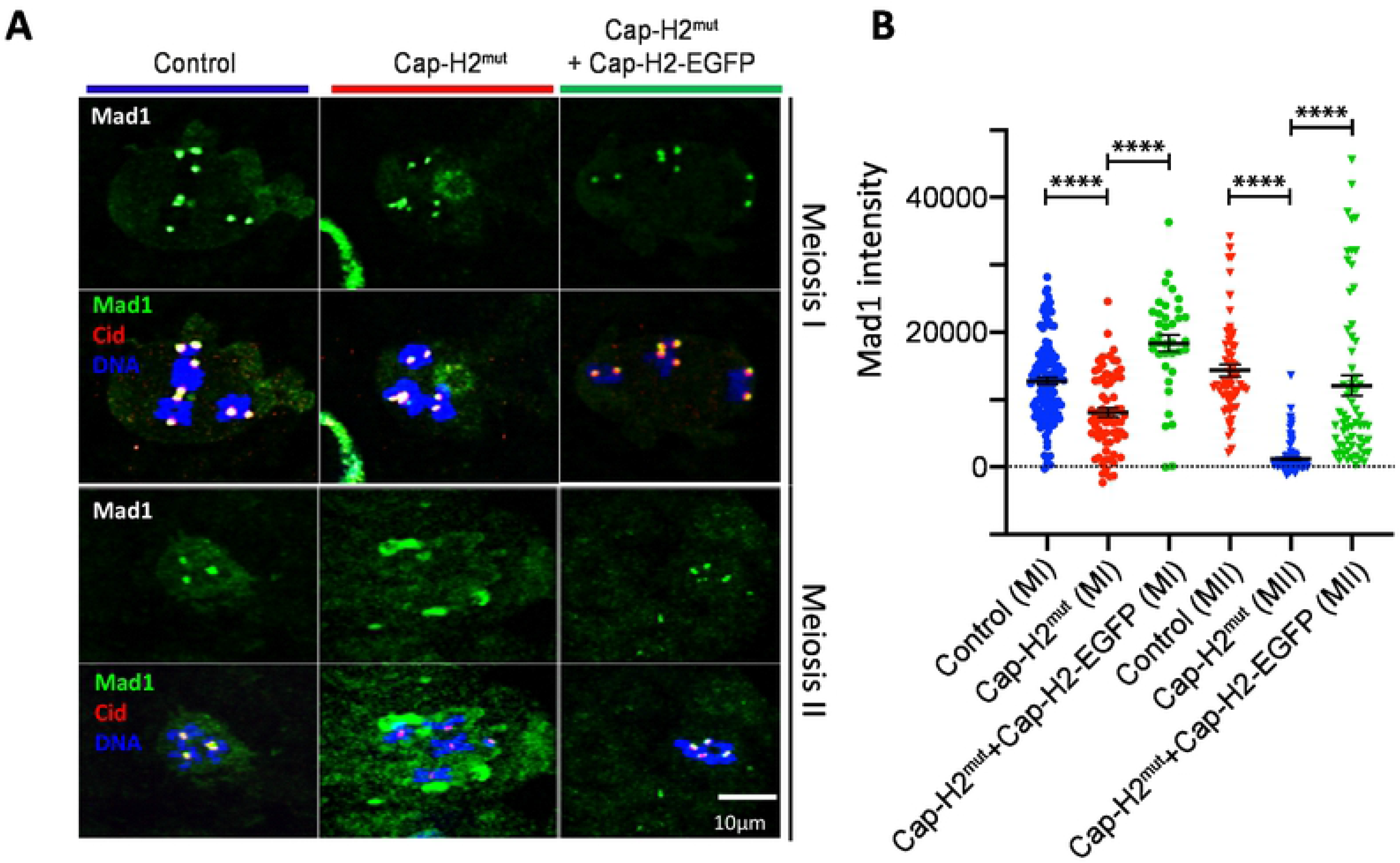
CAP-H2 mutants are less able to recruit the SAC Component Mad1 to the kinetochore. (**A**) Immunofluorescence of Mad1 protein (green) upon microtubules depolymerisation with colchicine treatment (100 μM) in meiosis I and meiosis II spermatocytes in Control, Cap-H2^mut^, and rescued strain (Cap-H2^mut^ + CapH2-EGFP). Cid (red) was also used as a centromere marker and DNA (blue) counterstained with DAPI. (**B**) Quantification of the Mad1 levels at each kinetochore, after background subtraction, at individualized unattached kinetochores after treatment with 100 μM colchicine. Each dot represents the average of all kinetochores per cell (8 co-joined kinetochore pairs (MI) or 8 single kinetochores (MII)), derived from at least 4 independent experiments. ****, P < 0.0001 using ONE-way ANOVA (“Tukey’s multiple comparisons test”). Black lines indicate Mean ± SEM. Detailed genotypes can be found in Table S3.

The levels of Mad1 at the nuclear periphery of interphase cells or its cytoplasmic pool during meiosis are not consistently decreased in Cap-H2 mutants (Fig. S3). These findings suggest that the reduced Mad1 amount at kinetochores stems from a defect in its recruitment and not from reduced protein levels. Importantly, this defect seems to be specific to meiotic divisions as mitotically dividing cells display normal levels of Mad1 upon microtubule depolymerization, both in mitotic spermatocytes and larval neuroblasts (Fig. S4). Thus, the inability to recruit Mad1 associated with Cap-H2 mutations, and likely to establish of a robust SAC response, is specific to meiotic divisions.

### Cap-H2 mutants have normal levels of centromere and kinetochore structural components

To access how chromatin structural problems could impair SAC activation, we aimed to access whether or not condensin mutants display defects in the recruitment of centromere and kinetochore structural components. In Hela cells and *Xenopus* egg extract, Condensin II was found to cooperate with Hjurp to the loading of the centromere marker CenpA, and thereby control centromeric chromatin [39, 40]. Additionally, a subpopulation of condensin I/II was reported to localize to the inner kinetochore plate in *C. elegans* and human cells [41, 50]. These studies would support a model whereby Condensin II could impair SAC response by impairing centromere/kinetochore organization. To test this notion, we compared the levels of several centromere and kinetochore components in both control and CapH2^mut^ cells using both immunofluorescence analysis and live cell imaging approaches. However, our results suggest that Condensin II mutants seem to have a normal kinetochore structure, at least at the level of resolution tested, as no major differences could be found in the relative amounts of Cenp-A/Cid, Cenp-C, Spc105 and Mis12, when comparing control and CAP-H2 mutant cells (Figure 4). Although we cannot exclude that condensin II may impose mild changes in kinetochore organization, not detectable in our analysis, we conclude that failures to recruit checkpoint components is independent of the recruitment of centromere/kinetochore core components, at least at the quantitative level.

**Figure 4.**
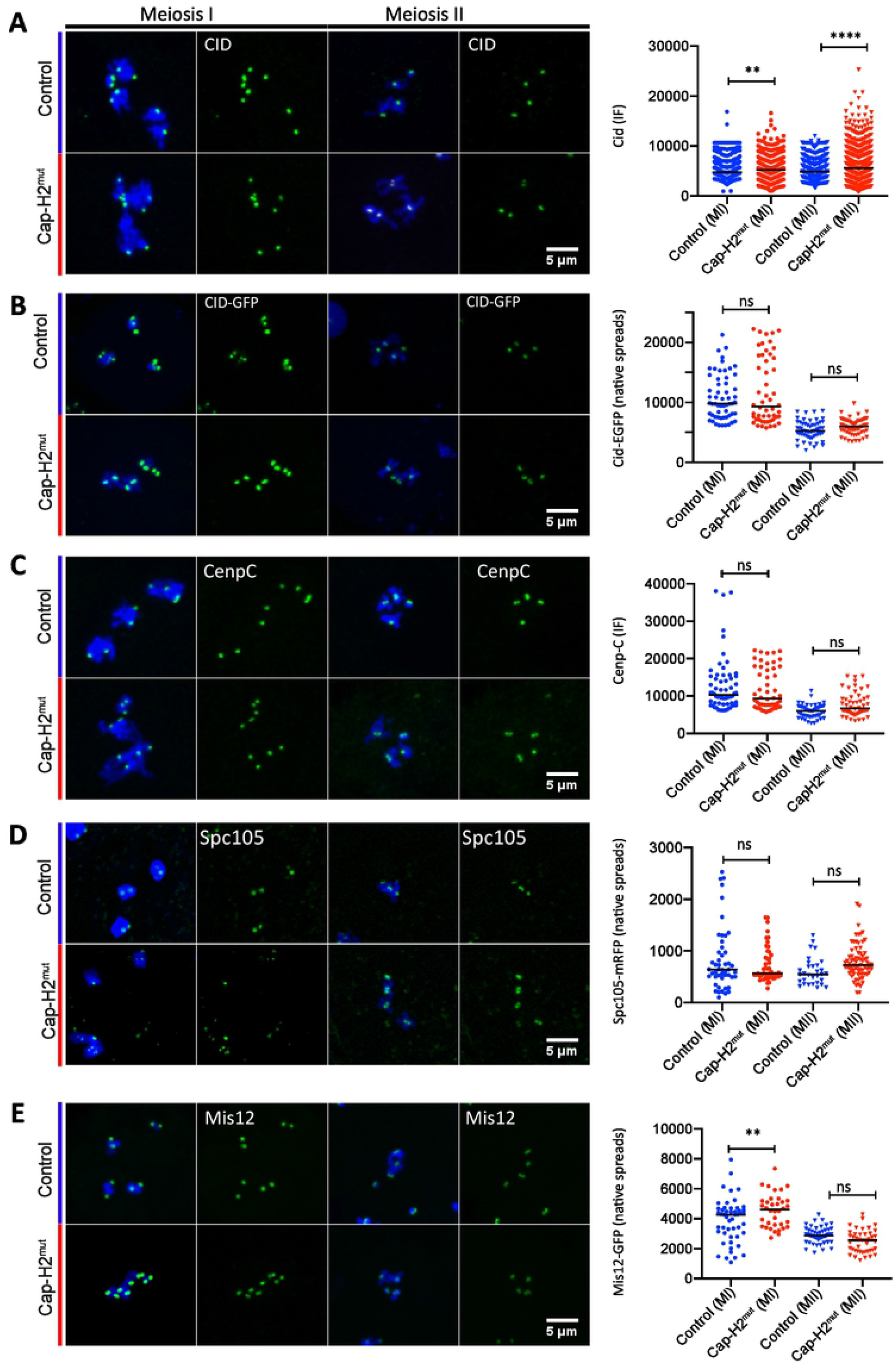
CAP-H2 mutant presents normal levels of centromere/kinetochore proteins. Representative images of immunostaining (**A, C**) or native testis spreads (**B, D, E**) of the following centromere/kinetochore proteins: Cid/CenpA (**A**and **B**); Cenp C (**C**) Spc105 (**D**) and Mis12 (**E**) in Control and Cap-H2^mut^ spermatocytes; Scale bars are 5μm and apply to all images; Right panels represent quantification of the fluorescence intensity for each analyzed protein. Each dot represents one kinetochore (A) or one cell (average of the signal measured in 8 centromeres/kinetochores) (B-E), derived from at least 4 independent experiments. *, P < 0.05; **, P < 0.01; ***, P < 0.001; ****, P < 0.0001 using ONE-way ANOVA (“Tukey’s multiple comparisons test”). Black lines indicate Median. Detailed genotypes can be found in Table S3.

### Reduced meiotic timing is the main driver of segregation errors observed in Condensin II mutants

Our results so far uncover a new role for condensin II in the efficiency of SAC response. We next aimed to test how this novel function contributes to the meiotic defects associated with Cap-H2 mutants. SAC-deficient males are reported to be fertile [46] and hence shortening of meiotic timing should not have an adverse effect on its own. However, we reasoned that in the context of condensin II impairment, short meiotic timings could limit the action of other components for chromosome organization, that would become rate-limiting in the absence of condensin II. Although both Condensin complexes have been proposed to have non-overlapping functions in chromosome organization in mammalian cells [2, 22], the fact that in *Drosophila* mitotic divisions rely mostly on Condensin I suggests that this complex is sufficient to fulfil the requirements for chromosome organization in this organism. Interestingly, a recent report highlights that condensin I in male meiosis is only loaded after nuclear envelop breakdown [23], in contrast to mitotic divisions, where Condensin I is already present on prophase chromosomes [15]. This finding further raises the hypothesis that the short meiotic timing observed in Cap-H2 may be limiting the action of Condensin I (now limited to the short time between NEBD and anaphase), which could preclude any compensation that Condensin I could have on meiotic chromosome assembly.

We therefore hypothesized that the accelerated meiosis timing could be limiting the action of other components involved in chromosome architecture. To test this hypothesis, we asked if imposing an artificial delay in meiosis progression (by bypassing SAC insufficiency) could improve the segregation errors observed in Cap-H2 mutant. First, we probed for conditions that could impose an artificial SAC-independent delay in meiotic progression. Our strategy was to lower the levels of APC/c, and therefore delay meiotic exit, using experimental conditions in which meiosis would last longer without impaired segregation fidelity. To probe for such conditions, we took advantage of the fact that APC/c is composed of several subunits to test several RNAi lines for some individual components (Table S1). Using Bam-Gal4 to specifically drive de RNAi expression in testis, we initially screened several APC/c subunits RNAi lines for male fertility, aiming to exclude RNAi lines that would cause such a strong APC/c impairment that would result in male sterility. For that, males expressing different RNAi lines under the control of Bam-Gal4 driver were crossed with wild-type females. The crosses were kept at 25°C and we accessed for the presence of progeny after two weeks. As expected, several of the lines tested were male sterile. However, 3 lines were found to be fertile and selected for further analysis. Live-cell imaging of selected individual lines was performed in order to estimate the division timing (Table S1). From all the lines tested, we selected the second chromosome insertion of Cdc23 RNAi line which presents a significant meiotic extension (~89 min compared to ~36 min in controls) (Figure 5 and Table S1).

**Figure 5.**
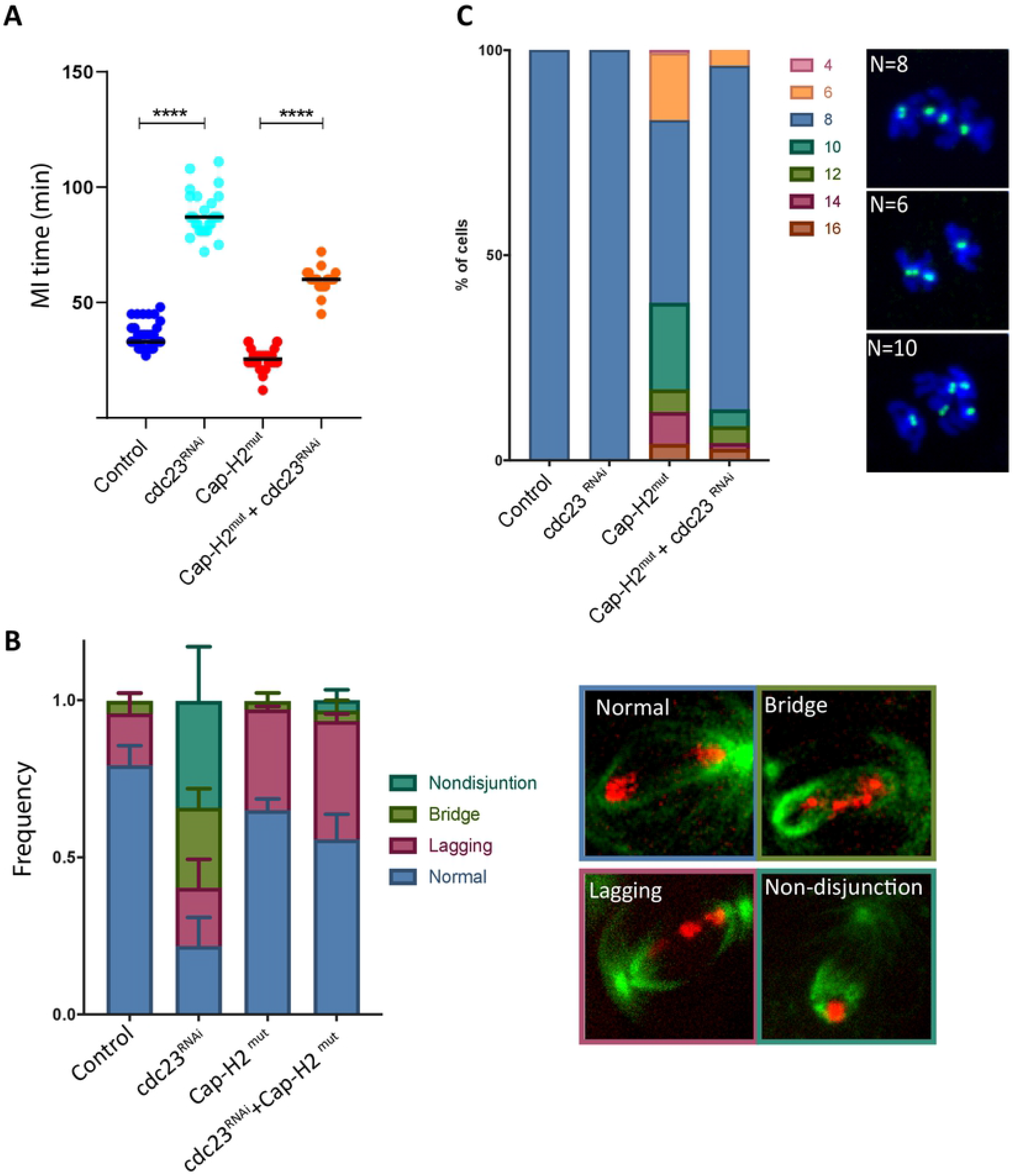
Reduced meiotic timing is the main driver of segregation errors observed in CAP-H2 mutants: (**A**) Timing of meiosis I division in Control, Cap-H2^mut^, cdc23RNAi and Cap-H2^mut^ + cdc23RNAi spermatocytes. *, P < 0.05; **, P < 0.01; ***, P < 0.001; ****, P < 0.0001 using ONE-way ANOVA (“Tukey’s multiple comparisons test”). Black lines indicate Median. **B**) Quantification of segregation errors observed at meiosis I exit in Control, Cap-H2^mut^, cdc23RNAi and Cap-H2^mut^+cdc23RNAi spermatocytes. Images depict representative pictures of the different error classes. Errors were quantified from at least 30 cells from > 3 independent movies. *, P < 0.05; **, P < 0.01; ***, P < 0.001; ****, P < 0.0001 using ONE-way ANOVA (“Tukey’s multiple comparisons test”). Black lines indicate Median. (**C**) Quantification of aneuploidy in meiosis II-arrested cells in the same conditions. The number of centromeres was scored manually using CID/CenpA staining. At least 50 cells were analysed for each condition. Detailed genotypes can be found in Table S3.

Next, we developed two assays to monitor the fidelity of meiosis I. The first one relies on the analysis of chromosome segregation defects by live cell imaging. The second is based on the analysis of chromosome number (based in CID/CenpA dots) present in cells arrested in meiosis II. This second method is an indirect read-out of the segregation defects in meiosis I as severe segregation defects should result in non-disjunction and abnormal chromosome numbers. As expected, CAP-H2 mutant displays a high degree of segregation errors (mostly chromatin bridges) and have a high aneuploidy rate in meiosis II spermatocytes, as a consequence of a catastrophic MI division (Figure 5). Remarkably, RNAi for Cdc23, despite the extended meiotic timing, did not perturb the fidelity of meiotic division, as shown by the reduced frequency of mitotic defects in MI and nearly absent levels of aneuploidy in MII (Figure 5). We therefore concluded that this experimental approach was able to induce an artificial extension of meiotic timing without compromising meiotic divisions, and could then be used to test our hypothesis.

For that purpose, we combined Cdc23 RNAi and Cap-H2 mutations and asked if imposing more time to the cell during meiosis I would be sufficient to improve the meiotic fidelity. We observed that giving more time for meiotic progression (from 25±5 min to 60±6 min) is sufficient to reduce the frequency of segregation defects associated with Cap-H2 mutations (from 78% observed in Cap-H2 mutants to 44% in the Cap-H2 mutant combined with Cdc23 RNAi (Figure 5)). Most importantly, we were also able to restore the ploidy of the cell and the frequency of cells presenting in average eight Cid/CenpA dots (corresponding to four chromosomes, the correct haploid number) is now 83%, compared to 45% observed in CAP-H2 mutants.

We therefore conclude that the short meiotic timing observed in condensin II mutants, due to its role in establishing a robust SAC response, is a major contributor of the meiotic defects associated with condensin II loss. Yet, rescue of meiotic fidelity is insufficient to restore male fertility as despite the improved fidelity in meiotic divisions, Cap-H2 males, combined with cdc23 RNAi remained sterile (data not shown). These findings imply that condensin II holds other unknown roles during spermatogenesis, beyond its well-established role in meiotic segregation.

## Discussion

One of the major phenotypes observed in cells deficient in condensin complexes is the massive amount of DNA bridges during anaphase in both meiosis and mitosis. These segregations defects have been mostly attributed to abnormal condensation or sister chromatid resolution problems.

Here we show, that in addition to the establishment of proper chromatin structure in Drosophila male meiosis, Condensin II (Cap-H2 and Cap-D3) is required for a robust SAC response. Furthermore, we demonstrate that upon Cap-H2 loss, the inability of prolonging meiosis is a major driver for the observed segregation defects and consequently aneuploidy observed upon Condensin II mutations.

It remains unknown how general this novel function is across the tree of life. Previous studies in mouse oocytes indicate a prolonged meiotic arrest in Cap-H2 mutants [9] arguing for a functional SAC in female meiosis in this system. Conversely, yeast strains carrying point mutations in condensin subunits are hypersensitive to spindle poisons which was proposed to be caused by insufficient SAC response [51]. *Drosophila* male meiosis is quite unique as bivalents are formed in the absence of recombination. It is therefore conceivable that the bivalent conformation may render these cells more dependent on Condensin II for robust SAC function. Moreover, other Condensin’s functions have been previously demonstrated to vary among different organisms, which may account for some organism or even tissue specificities.

Although we cannot exclude that accelerated meiotic timings may partly derive from abnormal kinetochore-microtubule attachments, previously suggested to depend on condensin-II [41, 49], we favour that condensin II has a direct role in the establishment of a robust SAC response, independently of MT attachments. This is mostly supported by the fact that CAP-H2 cells are less efficient in meiotic arrest when in the presence of microtubule poisons and display reduced levels of a critical SAC component (Mad1) at their kinetochores.

Failure to recruit Mad1 could be attributed to defects in kinetochore/centromere organization, as previously reported for condensin II in other systems [39–41, 50]. However, our results suggest that Condensin II mutants seem to have a normal kinetochore structure, at least at the level of resolution tested. Other mechanisms may therefore ensure efficient Mad1 recruitment, independently of centromere/kinetochore organization.

Regardless of the exact mode of action, our work highlights that this novel role for Condensin II in SAC robustness is a major contributor to the segregation defects observed in the mutants, since defects can be rescued by artificial prolonging meiotic duration. The real nature behind this “time-rescue” is not completely understood. We nevertheless favour that the observed rescue could be related to the presence of a second Condensin complex in the system. Giving more time to the cell could allow Condensin I to compensate the function of Condensin II. In *Drosophila* spermatocytes, Condensin I nuclear localization occurs only after NEBD [23], in contrast to what was previously reported for mitotic tissues, in which Condensin I localizes to the nucleus before NEBD [15]. In fact, the condensin I localization pattern in Drosophila male meiosis is closer to what has been described for mammalian cells where condensin II is the prime complex involved in prophase compaction while condensin I only acts after NEBD [7, 52]. This delayed chromatin access may restrict the action of Condensin I to a short time-period and hinder its ability to compensate for Condensin II deficiency. Therefore, by prolonging meiotic division, we give more access and time for Condensin I to act on meiotic chromatin and potentially to overcome the topological problems that results from Condensin II loss.

Intriguingly, even though we could rescue the segregation defects by prolonging meiotic division, this segregation rescue did not restore the fertility in those males. Our data is in line with previous reports where inhibition of autosome association was shown to rescue anaphase I bridges observed in condensin II mutants but is also insufficient to revert the associated sterility [18]. Taken together, these findings strongly argue that Condensin II may be required in a non-meiotic stage of spermatogenesis.

In summary, our results point that Condensin II is a major regulator *Drosophila* spermatogenesis, acting at multiple stages to ensure meiotic fidelity. In addition to the previously reported role in territories formation and chromatin resolution [18, 34], we provide evidence that Condensin II also contributes to the fidelity of segregation through the regulation of SAC response.

## Materials and Methods

### Fly Strains

*Drosophila melanogaster* flies were raised at 25°C or 18°C in polypropylene vials (51 mm diameter) containing enriched medium (cornmeal, molasses, yeast, soya flour and beetroot syrup).\ For CAP-H2 mutant conditions we used a CAP-H2 mutant allele containing a PBac(RB) insertion at the end of the third exon obtained from the Exelixis collection (CAP-H2e03210, see Fig. S1) over a deficiency for the same chromosomal region (Df(3R)BSC529). For live cell imaging, flies also expressed RFP-tagged Histone (H2Av-mRFP) [53], beta-tubulin tagged with EGFP (gift from Ivo Telley) and/or Cid/CenpA tagged with a EGFP [53]. Additional flies include: Mad2 mutant flies (mad2P, [46]); CAP-D3 mutant and rescued strain ([18] and unpublished strain by Evelin Urban and Stefan Heidmann); Mis12-EGFP [54]; Spc105-RFP [55]. For the APC/c RNAi lines screened, we used stocks from the Transgenic RNAi project (TRiP) available in the Bloomington Drosophila Stock Center. RNAi in the germiline was driven using Bam-Gal4 (from Michael Buszczak Lab).

For the rescue of Cap-H2 function, flies carrying an ectopic copy of the CAP-H2 gene and regulatory regions were produced by random P-element integration (Bestgene). Cap-H2 genomic region was amplified using 5’ GCATGAGCGGCCGCGGCGAATCACTCACGATAGTG 3’ and reverse 5’ GCATGAGGTACCCACAAGAACATGTGGGAGCTC 3 primers. The region was modified to include a EGFP-tag at C-terminus and include three consecutive TEV cleavage sites at position aa 398 (see details in Fig. S1).

A complete list of all the stocks and the specific genotypes used in each figure can be found in the supplemanty Tables S2 and S3, respectively.

### RT-PCR

RNA extraction of ovaries was performed using a Trizol based kit (Invitrogen) according to the manufacturer’s instructions. DNA contamination was removed by adding a DNase I (ThermoFisher Scientific) incubation step to the protocol. Reverse transcriptase Kit (ThermoFisher Scientific) was used to synthesize cDNA from the previous extracted RNA. The expression of CAP-H2 was tested by using the follow primers: forward 5’CCCAACCTGACAATTCAGACC 3’ and reverse 5‘CATCGAAGATATCCCCGAGT3’. For loading control Oscar 5‘-TTCCTCCAGCTGCCTCAAC-3’ and reverse 5’-GTTCGACTCCGCTGCCTCAT-3’ primers where used.

The primers was designed within an exon region to excluse a possible genomic DNA contamination fragment sizes. The final product was run in a 1.5% agarose gel with RedSafe (INtRON Biotechnology).

### Pupae testis Live Imaging

Testes were dissected from *Drosophila* pupae of the different strains in 1xPBS. After dissection, testes were placed in a 200μl drop of Schneider’s medium supplemented with 10% FBS (+ 100μM colchicine, when referred) on a glass-bottom petri dish (MakTek). Samples were imaged on a Leica spinning disc (Leica Microsystems, Germany) using a 63× (glycerol-immersion) objective. A total of 38 *Z*-stacks with a Z-step of 0.8μm was taken every 3 min. All the images were processed using FIJI software.

### Cytological analysis and Immunofluorescence

For immunofluorescence analysis of meiotic and premeiotic spermatocytes, pupae testes were dissected in PBS and incubated with Schneider’s medium supplemented with 10%FBS and 100μM colchicine for 45minutes. After this period, testes were incubated in a hypotonic solution of sodium citrate 0.5% for 2 min and fixed in PBS-0.05%Tween with 3.7% formaldehyde for 30sec-1min. Testes were transferred onto a siliconized coverslip and squashed between a poly-L-lysine-coated slide and the coverslip. After that, slides were placed in liquid nitrogen and after coverslip removal, immunofluorescence protocol was performed on the slide. Testes were washed with 1× PBS and extracted for 10 min in 1× PBS with 0.1% Triton X-100 with gentle agitation. Slides were then washed 3x 5min in 0.05%PBS-Tween 20. Unspecific binding of primary antibodies was blocked by incubating samples with 0.05% Tween, 5% FBS PBS for 30min.

For brain spreads and immunofluorescence, third-instar larvae brains were dissected in PBS, incubated with 100 μM colchicine for one hour, hypotonic shocked in 0.5% sodium citrate for 2–3 min, and fixed on a 5-μl drop of fixative (3.7% formaldehyde, 0.1% Triton-X100 in PBS) placed on top of a siliconized coverslip. After 30 seconds, the brains were squashed between the coverslip and a slide, allowed to fix for an additional 1 min, and then placed in liquid nitrogen. Slides were further extracted with 0.1% Triton X-100 in PBS for 10 min and used for immunofluorescence following standard protocols.

In both cases, primary antibodies were diluted in the blocking solution and incubated overnight. Used dilutions were as follows: rat anti-CID/CenpA, 1:5000 (gift from C.E. Sunkel), rabbit anti-Mad1, 1:1000 [56] and rabbit anti-CENPC, 1:5000 [57]. All secondary antibodies were from Alexa Fluor (Thermo Scientific) and were used at 1:500 dilution. Finally, after the wash of secondary antibodies, samples were mounted in Vectashield with DAPI (Vector Laboratories, Burlingame, CA, USA).

For nurse cells morphology imaging, females of each genotype were maintained for 2 days in Vienna food with additional yeast. The ovaries were then dissected in PBS and fixed in a 1:3 solution of PBX* (PBS, 1%, 10% NP-40, 4% formaldehyde) and heptane for 20 minutes with gentle agitation. The ovaries were then washed in PBST (PBS and 0.2% Tween) three times for 5 minutes and permeabilized for 1 hour in PBST*(PBST plus 1% Triton). DNA was stained with DAPI present in the VECTASHIELD mounting medium (VectorLabs) or by an additional 20 min incubation step with Sytox green (ThermoFisher Scientific) solution (PBST plus 0.02% Sytox green, and 1%RNAase).

To probe for the localization of centromeric/kinetochore proteins, GFP/RFP versions of the proteins were accessed using non-fixed (native) chromosome spreads. After incubation with 100μM colchicine, 6-8 testes were placed in a 15μl PBS supplemented with 100μM colchicine and 2 μg/ml Hoechst 33258 and immediately squashed between a slide and a coverslip by capillary forces. Images were taken up to 30min after sample preparation. Fluorescence images were acquired on a Leica spinning disc (Leica Microsystems, Germany) using a 63× (glycerol-immersion) objective.

### Quantitative imaging analysis

For analysis of meiotic timing, we monitored the time between NEBD (defined as by the time soluble histones are dispersed) to anaphase onset (defined by chromosome movement towards opposite poles). In colchicine arrest experiments, meiotic exit was defined by decondensation of chromatin. Segregation errors were scored manually, based on HisH2Av signal.

For the quantification of immunofluorescence images to probe for Cid/Mad1 levels at centromeres/kinetochores, we used Z-projections of each cell (made in FIJI), and a manually placed fixed-size ROI (7μm for MI and 5μm for MII) was used to select for the centromere/kinetochore area, based on Cid/CenpA signal. The mean intensity of Cid/Mad1 signal within that region was measured. To estimate for cytoplasmic background levels, a separate ROI was used, placed next to the centromeres and used to subtract to each centromere signal. For the quantification of centromere/kinetochore proteins using native chromosome spreads, images were background subtracted and segmented to select the chromosomal area (based on Hoechst signal). For each metaphase, an automatic default threshold was used within each area to select for centromeres/kinetochore signals and the mean fluorescence intensity of all centromeres/kinetochores was measured using FIJI software.

### Fertility tests

For each genotype tested, individual newly ecloded males (max 3-day-old) were crossed with wild type virgin females (Oregon-R). For the APC screening males from the tested UAS-RNAi lines were crossed with females carrying a Bam-Gal4 driver. Individual newly ecloded F1 males (max 3-day-old) from each cross (expressing RNAi for the for different APC/c components in the germline) were then crossed with wild type (w*) virgins and kept at 25°C for two weeks. The presence of progeny (larvae and pupae) was determined after this period in both cases. For each genotype, a total of four replicates were made.

## Acknowledgements

We thank S. Heidmann, R. Karess, C. Lehner, C.E. Sunkel, M. Buszczak, I. Telley Drosophila Bloomington Stock Center (DBSC) and the Exelixis Drosophila Collection for fly strains and antibodies, and all the members of the R.A. Oliveira laboratory for discussions and comments. We thank the technical support of Instituto Gulbenkian de Ciencia’s Advanced Imaging Facility, supported by national Portuguese funding (PPBI-POCI-01-0145-FEDER-022122), and the Fly Facility, supported by Congento (LISBOA-01-0145-FEDER-022170). These two programs are cofinanced by Lisboa Regional Operational Program (Lisboa 2020) under the Portugal 2020 Partnership Agreement through the European Regional Development Fund (FEDER) and Fundação para a Ciência e a Tecnologia (FCT; Portugal) This work had the following financial support: doctoral fellowship by Fundação para a Ciência e Tecnologia (FCT) awarded to CH (SFRH/BD/113758/2015); European Research Council (ERC) Starting Grant (2014-STG-638917) and EMBO Installation Grant (IG2778) to RAO.

## REFERENCES

1. Hirano T, Kobayashi R, Hirano M. Condensins, chromosome condensation protein complexes containing XCAP-C, XCAP-E and a Xenopus homolog of the Drosophila Barren protein. Cell. 1997;89(4):511–21. PubMed PMID: 9160743.

2. Ono T, Losada A, Hirano M, Myers MP, Neuwald AF, Hirano T. Differential contributions of condensin I and condensin II to mitotic chromosome architecture in vertebrate cells. Cell. 2003;115(1):109–21. PubMed PMID: 14532007.

3. Yatskevich S, Rhodes J, Nasmyth K. Organization of Chromosomal DNA by SMC Complexes. Annu Rev Genet. 2019;53:445–82. Epub 2019/10/03. doi: 10.1146/annurev-genet-112618-043633. PubMed PMID: 31577909.

4. Ganji M, Shaltiel IA, Bisht S, Kim E, Kalichava A, Haering CH, et al. Real-time imaging of DNA loop extrusion by condensin. Science. 2018;360(6384):102–5. Epub 2018/02/24. doi: 10.1126/science.aar7831. PubMed PMID: 29472443; PubMed Central PMCID: PMCPMC6329450.

5. Kim E, Kerssemakers J, Shaltiel IA, Haering CH, Dekker C. DNA-loop extruding condensin complexes can traverse one another. Nature. 2020;579(7799):438–42. Epub 2020/03/07. doi: 10.1038/s41586-020-2067-5. PubMed PMID: 32132705.

6. Oliveira RA, Coelho PA, Sunkel CE. The condensin I subunit Barren/CAP-H is essential for the structural integrity of centromeric heterochromatin during mitosis. Mol Cell Biol. 2005;25(20):8971–84. PubMed PMID: 16199875.

7. Gerlich D, Hirota T, Koch B, Peters JM, Ellenberg J. Condensin I Stabilizes Chromosomes Mechanically through a Dynamic Interaction in Live Cells. Current biology : CB. 2006;16(4):333–44. PubMed PMID: 16488867.

8. Ribeiro SA, Gatlin JC, Dong Y, Joglekar A, Cameron L, Hudson DF, et al. Condensin regulates the stiffness of vertebrate centromeres. Mol Biol Cell. 2009;20(9):2371–80. Epub 2009/03/06. doi: E08-11-1127 [pii] 10.1091/mbc.E08-11-1127. PubMed PMID: 19261808.

9. Houlard M, Godwin J, Metson J, Lee J, Hirano T, Nasmyth K. Condensin confers the longitudinal rigidity of chromosomes. Nature cell biology. 2015;17(6):771–81. doi: 10.1038/ncb3167. PubMed PMID: 25961503.

10. Piskadlo E, Oliveira RA. A Topology-Centric View on Mitotic Chromosome Architecture. Int J Mol Sci. 2017;18(12). doi: 10.3390/ijms18122751. PubMed PMID: 29258269; PubMed Central PMCID: PMCPMC5751350.

11. Piskadlo E, Tavares A, Oliveira RA. Metaphase chromosome structure is dynamically maintained by condensin I-directed DNA (de)catenation. Elife. 2017;6. doi: 10.7554/eLife.26120. PubMed PMID: 28477406; PubMed Central PMCID: PMCPMC5451211.

12. Shintomi K, Hirano T. The relative ratio of condensin I to II determines chromosome shapes. Genes Dev. 2011;25(14):1464–9. Epub 2011/07/01. doi: 10.1101/gad.2060311. PubMed PMID: 21715560; PubMed Central PMCID: PMCPMC3143936.

13. Hoencamp C, Dudchenko O, Elbatsh AMO, Brahmachari S, Raaijmakers JA, van Schaik T, et al. 3D genomics across the tree of life reveals condensin II as a determinant of architecture type. Science. 2021;372(6545):984–9. Epub 2021/05/29. doi: 10.1126/science.abe2218. PubMed PMID: 34045355; PubMed Central PMCID: PMCPMC8172041.

14. Herzog S, Nagarkar Jaiswal S, Urban E, Riemer A, Fischer S, Heidmann SK. Functional dissection of the Drosophila melanogaster condensin subunit Cap-G reveals its exclusive association with condensin I. PLoS genetics. 2013;9(4):e1003463. doi: 10.1371/journal.pgen.1003463. PubMed PMID: 23637630; PubMed Central PMCID: PMCPMC3630105.

15. Oliveira RA, Heidmann S, Sunkel CE. Condensin I binds chromatin early in prophase and displays a highly dynamic association with Drosophila mitotic chromosomes. Chromosoma. 2007;116(3):259–74. Epub 2007/02/24. doi: 10.1007/s00412-007-0097-5. PubMed PMID: 17318635.

16. Nguyen HQ, Nye J, Buster DW, Klebba JE, Rogers GC, Bosco G. Drosophila casein kinase I alpha regulates homolog pairing and genome organization by modulating condensin II subunit Cap-H2 levels. PLoS genetics. 2015;11(2):e1005014. Epub 2015/02/28. doi: 10.1371/journal.pgen.1005014. PubMed PMID: 25723539; PubMed Central PMCID: PMCPMC4344196.

17. Buster DW, Daniel SG, Nguyen HQ, Windler SL, Skwarek LC, Peterson M, et al. SCFSlimb ubiquitin ligase suppresses condensin II-mediated nuclear reorganization by degrading Cap-H2. The Journal of cell biology. 2013;201(1):49–63. doi: 10.1083/jcb.201207183. PubMed PMID: 23530065; PubMed Central PMCID: PMCPMC3613687.

18. Hartl TA, Sweeney SJ, Knepler PJ, Bosco G. Condensin II resolves chromosomal associations to enable anaphase I segregation in Drosophila male meiosis. PLoS genetics. 2008;4(10):e1000228. doi: 10.1371/journal.pgen.1000228. PubMed PMID: 18927632; PubMed Central PMCID: PMCPMC2562520.

19. Hartl TA, Smith HF, Bosco G. Chromosome alignment and transvection are antagonized by condensin II. Science. 2008;322(5906):1384–7. Epub 2008/11/29. doi: 322/5906/1384 [pii] 10.1126/science.1164216. PubMed PMID: 19039137.

20. Condensins: Universal organizers of chromosomes with diverse functions, (2012).

21. Yu HG, Koshland D. Chromosome morphogenesis: condensin-dependent cohesin removal during meiosis. Cell. 2005;123(3):397–407. PubMed PMID: 16269332.

22. Lee J, Ogushi S, Saitou M, Hirano T. Condensins I and II are essential for construction of bivalent chromosomes in mouse oocytes. Mol Biol Cell. 2011;22(18):3465–77. Epub 2011/07/29. doi: 10.1091/mbc.E11-05-0423. PubMed PMID: 21795393; PubMed Central PMCID: PMCPMC3172270.

23. Kleinschnitz K, Viessmann N, Jordan M, Heidmann SK. Condensin I is required for faithful meiosis in Drosophila males. Chromosoma. 2020;129(2):141–60. Epub 2020/04/22. doi: 10.1007/s00412-020-00733-w. PubMed PMID: 32314039; PubMed Central PMCID: PMCPMC7260282.

24. Hagstrom KA, Holmes VF, Cozzarelli NR, Meyer BJ. C. elegans condensin promotes mitotic chromosome architecture, centromere organization, and sister chromatid segregation during mitosis and meiosis. Genes Dev. 2002;16(6):729–42. PubMed PMID: 11914278.

25. Chan RC, Severson AF, Meyer BJ. Condensin restructures chromosomes in preparation for meiotic divisions. The Journal of cell biology. 2004;167(4):613–25. PubMed PMID: 15557118.

26. Smith SJ, Osman K, Franklin FC. The condensin complexes play distinct roles to ensure normal chromosome morphogenesis during meiotic division in Arabidopsis. Plant J. 2014;80(2):255–68. Epub 2014/07/30. doi: 10.1111/tpj.12628. PubMed PMID: 25065716; PubMed Central PMCID: PMCPMC4552968.

27. Siddiqui NU, Stronghill PE, Dengler RE, Hasenkampf CA, Riggs CD. Mutations in Arabidopsis condensin genes disrupt embryogenesis, meristem organization and segregation of homologous chromosomes during meiosis. Development. 2003;130(14):3283–95. Epub 2003/06/05. doi: 10.1242/dev.00542. PubMed PMID: 12783798.

28. Howard-Till R, Loidl J. Condensins promote chromosome individualization and segregation during mitosis, meiosis, and amitosis in Tetrahymena thermophila. Mol Biol Cell. 2018;29(4):466–78. Epub 2017/12/15. doi: 10.1091/mbc.E17-07-0451. PubMed PMID: 29237819; PubMed Central PMCID: PMCPMC6014175.

29. Yu HG, Koshland DE. Meiotic condensin is required for proper chromosome compaction, SC assembly, and resolution of recombination-dependent chromosome linkages. The Journal of cell biology. 2003;163(5):937–47. PubMed PMID: 14662740.

30. Hong S, Choi EH, Kim KP. Ycs4 is Required for Efficient Double-Strand Break Formation and Homologous Recombination During Meiosis. J Microbiol Biotechnol. 2015;25(7):1026–35. Epub 2015/05/16. doi: 10.4014/jmb.1504.04013. PubMed PMID: 25975613.

31. Li P, Jin H, Yu HG. Condensin suppresses recombination and regulates double-strand break processing at the repetitive ribosomal DNA array to ensure proper chromosome segregation during meiosis in budding yeast. Mol Biol Cell. 2014;25(19):2934–47. Epub 2014/08/12. doi: 10.1091/mbc.E14-05-0957. PubMed PMID: 25103240; PubMed Central PMCID: PMCPMC4230583.

32. Mets DG, Meyer BJ. Condensins regulate meiotic DNA break distribution, thus crossover frequency, by controlling chromosome structure. Cell. 2009;139(1):73–86. Epub 2009/09/29. doi: 10.1016/j.cell.2009.07.035. PubMed PMID: 19781752; PubMed Central PMCID: PMCPMC2785808.

33. Vazquez J, Belmont AS, Sedat JW. The dynamics of homologous chromosome pairing during male Drosophila meiosis. Current biology : CB. 2002;12(17):1473–83. Epub 2002/09/13. doi: 10.1016/s0960-9822(02)01090-4. PubMed PMID: 12225662.

34. Vernizzi L, Lehner CF. Bivalent individualization during chromosome territory formation in Drosophila spermatocytes by controlled condensin II protein activity and additional force generators. PLoS genetics. 2021;17(10):e1009870. Epub 2021/10/21. doi: 10.1371/journal.pgen.1009870. PubMed PMID: 34669718; PubMed Central PMCID: PMCPMC8559962.

35. Burrack LS, Applen Clancey SE, Chacon JM, Gardner MK, Berman J. Monopolin recruits condensin to organize centromere DNA and repetitive DNA sequences. Mol Biol Cell. 2013;24(18):2807–19. Epub 2013/07/26. doi: 10.1091/mbc.E13-05-0229. PubMed PMID: 23885115; PubMed Central PMCID: PMCPMC3771944.

36. Brito IL, Yu HG, Amon A. Condensins promote coorientation of sister chromatids during meiosis I in budding yeast. Genetics. 2010;185(1):55–64. Epub 2010/03/03. doi: 10.1534/genetics.110.115139. PubMed PMID: 20194961; PubMed Central PMCID: PMCPMC2870976.

37. Wassmann K. Sister chromatid segregation in meiosis II: deprotection through phosphorylation. Cell Cycle. 2013;12(9):1352–9. Epub 2013/04/12. doi: 10.4161/cc.24600. PubMed PMID: 23574717; PubMed Central PMCID: PMCPMC3674063.

38. Hernandez MR, Davis MB, Jiang J, Brouhard EA, Severson AF, Csankovszki G. Condensin I protects meiotic cohesin from WAPL-1 mediated removal. PLoS genetics. 2018;14(5):e1007382. Epub 2018/05/17. doi: 10.1371/journal.pgen.1007382. PubMed PMID: 29768402; PubMed Central PMCID: PMCPMC5973623.

39. Barnhart-Dailey MC, Trivedi P, Stukenberg PT, Foltz DR. HJURP interaction with the condensin II complex during G1 promotes CENP-A deposition. Mol Biol Cell. 2017;28(1):54–64. Epub 2016/11/04. doi: 10.1091/mbc.E15-12-0843. PubMed PMID: 27807043; PubMed Central PMCID: PMCPMC5221629.

40. Bernad R, Sanchez P, Rivera T, Rodriguez-Corsino M, Boyarchuk E, Vassias I, et al. Xenopus HJURP and condensin II are required for CENP-A assembly. The Journal of cell biology. 2011;192(4):569–82. Epub 2011/02/16. doi: 10.1083/jcb.201005136. PubMed PMID: 21321101; PubMed Central PMCID: PMCPMC3044122.

41. Samoshkin A, Arnaoutov A, Jansen LE, Ouspenski I, Dye L, Karpova T, et al. Human condensin function is essential for centromeric chromatin assembly and proper sister kinetochore orientation. PloS one. 2009;4(8):e6831. Epub 2009/08/29. doi: 10.1371/journal.pone.0006831. PubMed PMID: 19714251; PubMed Central PMCID: PMCPMC2730017.

42. Musacchio A. The Molecular Biology of Spindle Assembly Checkpoint Signaling Dynamics. Current biology : CB. 2015;25(20):R1002–18. Epub 2015/10/21. doi: 10.1016/j.cub.2015.08.051. PubMed PMID: 26485365.

43. Hirota T, Gerlich D, Koch B, Ellenberg J, Peters JM. Distinct functions of condensin I and II in mitotic chromosome assembly. Journal of cell science. 2004;117(Pt 26):6435–45. PubMed PMID: 15572404.

44. Watrin E, Legagneux V. Contribution of hCAP-D2, a non-SMC subunit of condensin I, to chromosome and chromosomal protein dynamics during mitosis. Mol Cell Biol. 2005;25(2):740–50. PubMed PMID: 15632074.

45. Uchida KS, Takagaki K, Kumada K, Hirayama Y, Noda T, Hirota T. Kinetochore stretching inactivates the spindle assembly checkpoint. The Journal of cell biology. 2009;184(3):383–90. doi: 10.1083/jcb.200811028. PubMed PMID: 19188492; PubMed Central PMCID: PMC2646554.

46. Buffin E, Emre D, Karess RE. Flies without a spindle checkpoint. Nature cell biology. 2007;9(5):565–72. Epub 2007/04/10. doi: ncb1570 [pii] 10.1038/ncb1570. PubMed PMID: 17417628.

47. Chaurasia S, Lehner CF. Dynamics and control of sister kinetochore behavior during the meiotic divisions in Drosophila spermatocytes. PLoS genetics. 2018;14(5):e1007372. Epub 2018/05/08. doi: 10.1371/journal.pgen.1007372. PubMed PMID: 29734336; PubMed Central PMCID: PMCPMC5957430.

48. Rebollo E, Gonzalez C. Visualizing the spindle checkpoint in Drosophila spermatocytes. EMBO reports. 2000;1(1):65–70. Epub 2001/03/21. doi: 10.1093/embo-reports/kvd011. PubMed PMID: 11256627; PubMed Central PMCID: PMCPMC1083687.

49. Wignall SM, Deehan R, Maresca TJ, Heald R. The condensin complex is required for proper spindle assembly and chromosome segregation in Xenopus egg extracts. The Journal of cell biology. 2003;161(6):1041–51. PubMed PMID: 12821643.

50. Maddox PS, Oegema K, Desai A, Cheeseman IM. “Holo”er than thou: chromosome segregation and kinetochore function in C. elegans. Chromosome Res. 2004;12(6):641–53. Epub 2004/08/04. doi: 10.1023/B:CHRO.0000036588.42225.2f. PubMed PMID: 15289669.

51. Xu X, Nakazawa N, Yanagida M. Condensin HEAT subunits required for DNA repair, kinetochore/centromere function and ploidy maintenance in fission yeast. PloS one. 2015;10(3):e0119347. Epub 2015/03/13. doi: 10.1371/journal.pone.0119347. PubMed PMID: 25764183; PubMed Central PMCID: PMCPMC4357468.

52. Ono T, Fang Y, Spector DL, Hirano T. Spatial and temporal regulation of Condensins I and II in mitotic chromosome assembly in human cells. Mol Biol Cell. 2004;15(7):3296–308. PubMed PMID: 15146063.

53. Schuh M, Lehner CF, Heidmann S. Incorporation of Drosophila CID/CENP-A and CENP-C into Centromeres during Early Embryonic Anaphase. Current biology : CB. 2007;17(3):237–43. PubMed PMID: 17222555.

54. Schittenhelm RB, Heeger S, Althoff F, Walter A, Heidmann S, Mechtler K, et al. Spatial organization of a ubiquitous eukaryotic kinetochore protein network in Drosophila chromosomes. Chromosoma. 2007;116(4):385–402. Epub 2007/03/03. doi: 10.1007/s00412-007-0103-y. PubMed PMID: 17333235.

55. Schittenhelm RB, Chaleckis R, Lehner CF. Intrakinetochore localization and essential functional domains of Drosophila Spc105. The EMBO journal. 2009;28(16):2374–86. Epub 2009/07/11. doi: emboj2009188 [pii] 10.1038/emboj.2009.188. PubMed PMID: 19590494.

56. Conde C, Osswald M, Barbosa J, Moutinho-Santos T, Pinheiro D, Guimaraes S, et al. Drosophila Polo regulates the spindle assembly checkpoint through Mps1-dependent BubR1 phosphorylation. The EMBO journal. 2013;32(12):1761–77. Epub 2013/05/21. doi: 10.1038/emboj.2013.109. PubMed PMID: 23685359; PubMed Central PMCID: PMCPMC3680734.

57. Heeger S, Leismann O, Schittenhelm R, Schraidt O, Heidmann S, Lehner CF. Genetic interactions of separase regulatory subunits reveal the diverged Drosophila Cenp-C homolog. Genes Dev. 2005;19(17):2041–53. Epub 2005/09/06. doi: 19/17/2041 [pii] 10.1101/gad.347805. PubMed PMID: 16140985.

